# Extended-depth of field random illumination microscopy, EDF-RIM, provides super-resolved projective imaging

**DOI:** 10.1101/2023.10.30.564754

**Authors:** Lorry Mazzella, Thomas Mangeat, Guillaume Giroussens, Benoit Rogez, Hao Li, Justine Creff, Mehdi Saadaoui, Carla Martins, Ronan Bouzignac, Simon Labouesse, Jérome Idier, Frédéric Galland, Marc Allain, Anne Sentenac, Loïc LeGoff

## Abstract

The ultimate aim of fluorescence microscopy is to achieve high-resolution imaging of increasingly larger biological samples. Extended depth of field presents a potential solution to accelerate imaging of large samples when compression of information along the optical axis is not detrimental to the interpretation of images. We have implemented an Extended Depth of Field (EDF) approach in a Random Illumination Microscope (RIM). RIM uses multiple speckled illuminations and variance data processing to double the resolution. It is particularly adapted to the imaging of thick samples as it does not require the knowledge of illumination patterns. We demonstrate highly-resolved projective images of biological tissues and cells. Compared to a sequential scan of the imaged volume with conventional 2D-RIM, EDF-RIM allows an order of magnitude improvement in speed and light dose reduction, with comparable resolution. As the axial information is lost in an EDF modality, we propose a method to retrieve the sample topography for samples that are organized in cell sheets.

## 1 Introduction

Fluorescence imaging has transformed biology by facilitating the observation of molecular events within live cells and tissues. One challenge is now to observe large samples (such as embryos) at the highest possible resolution^1^. Two major difficulties are faced when imaging large three-dimensional (3D) samples. First, the resolution over a wide field of view is challenged by optical aberrations. Second, imaging a large volume is slow, as it requires the sequential acquisition of many planes. It comes as a major challenge to simultaneously enhance spatio-temporal resolution and accommodate the imaging of increasingly larger samples as most live superresolution techniques tend to be slow and not adapted to thick samples.

Among the various methods for achieving super-resolution, structured illumination microscopy (SIM) presents a favorable balance between spatio-temporal resolutions and phototoxicity^2–8^. In SIM, the super-resolved image is computationally generated^9^ using multiple low-resolution images obtained with a periodic illumination pattern that is translated and rotated. SIM is amenable for live imaging and has been implemented in a number of modalities: light sheet microscopy^10–12^, total internal reflection microscopy^13–17^, 3D-microscopy^18,19^. However, SIM data processing requires the knowledge of the illumination pattern with great accuracy^20,21^, which makes the technique sensitive to aberrations and undermines its use in tissues. An active field of research, therefore, aims at improving the characterization of illumination patterns^9,20,22^, and to develop adaptive optics approaches^23,24^, or post-processing^25,26^, to correct potential artifacts. Another strategy was pursued in a variant of SIM called Random Illumination Microscopy (RIM). RIM replaces the periodic patterns with random speckled illuminations. It uses a reconstruction procedure based on the variance of the raw images which avoids the knowledge of the different illumination patterns. RIM has been demonstrated to be robust against aberrations and to achieve a two-fold improvement in lateral resolution compared to standard microscopy, while also providing strong sectioning in the axial direction^27^.

If some SIM approaches are becoming amenable to tissue imaging, the need to acquire multiple images of the sample under different illuminations at each observation plane still hinders the widespread use of SIM for large volume imaging. When the information from the biological sample can be compressed along the optical axis, extended depth of field (EDF) imaging becomes an interesting option for accelerating the imaging process through optical projection. Two main approaches are used for achieving extended depth of field. The first approach involves rapid scanning of the imaged plane with a tunable lens^28,29^, acoustic lens^30^ or a deformable mirror^31,32^. The second approach involves point spread function (PSF) engineering. Excitation PSF in scanning microscopes can be extended using, for example, bessel beams^33–36^. Detection PSF can also be elongated using phase masks^37^. EDF imaging is attractive because all the objects scattered along the optical axis appear in focus. On the other hand, because of the image projection, their footprint in focus is superimposed on all their defocused marks. As a result, EDF images are often affected by an important background, requiring debluring approaches^31,38^. This loss of contrast together with the distortion of the illumination pattern induced by the possibly thick sample has hampered the implementation of classical SIM in an extended depth of field configuration. As a result, we are not aware of any super-resolved extended depth microscope adapted to live biological samples.

Here, we combine a super-resolved technique with extended depth capacity using the RIM modality. The technique consists in illuminating the sample with multiple speckled illuminations and collecting light with a detection scheme featuring EDF capacity. The super-resolved image is formed from the raw images using the variance of the images and speckles. We demonstrate that RIM reconstruction process efficiently removes the background noise typically present in EDF images and improves the resolution of the deconvolved EDF widefield microscope by a factor of 1.7. We also demonstrate a strategy to partially compensate for the lost information along the optical axis through topographical estimation of the tissue on which the EDF images can be projected.

## 2 Theory

### 2.1 Introduction to Random illumination Microscopy (RIM)

Introducing *ρ* the sample fluorescence density, *h* the PSF of the optical system and *S* the illumination intensity, the intensity recorded at the camera plane of a fluorescence microscope (assuming a magnification of one) reads^39^

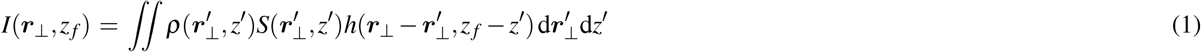

where *z* _*f*_ indicates the position of the focal plane, and the subscript ⊥ indicates transverse coordinates.

In Random Illumination Microscopy (RIM), a series of random speckle illuminations *S*_1_, · · · *S*_*M*_ are generated to provide a stack of speckled images. Each image has a resolution limit strictly enforced by the diffraction of light but the speckle illuminations, like the periodic light grid of SIM^2^, open a way toward super-resolution. It was demonstrated that a two-fold improvement in resolution compared to standard fluorescence microscopy could be achieved by processing the variance of the speckled images^27^. The key point of RIM is that the variance of the speckled images depends only on the observation point spread function and on the speckle auto-covariance, which are known functions. Up to now, RIM reconstruction algorithm has been implemented in the two-dimensional (2D) case^27^. The sample was assumed to be infinitely thin at the focal plane, 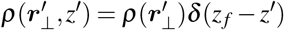 and the out-of-focus fluorescence reaching the camera was considered as a background noise^40^. The super-resolved reconstruction of the fluorescence density was achieved by estimating iteratively the sample so as to minimize the distance between the *experimental* standard deviation of the speckled images and its theoretical counterpart (see Methods for further details),

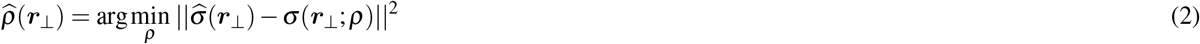

with 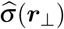 the experimental standard-deviation and *σ* (***r***_⊥_; *ρ*) the *theoretical* standard deviation derived from the expected variance that reads

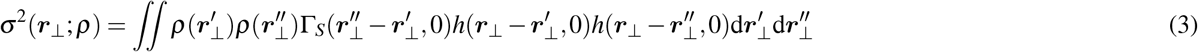

with 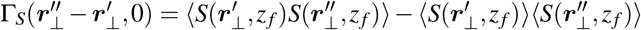 the auto-covariance of the speckle illuminations at the focal plane. Note that the 2D RIM reconstruction algorithm was able to form three-dimensional images of a sample by simply stacking 2D reconstructions obtained independently at the different focal planes.

### 2.2 Introduction to Extended Depth of Field imaging

For Extended-Depth of Field (EDF) imaging, it is necessary to consider *ρ, h*, and *S* as 3D functions. More precisely, the EDF image ∫ *I*_⊥_(***r***_⊥_) = ∫*I*(***r***_⊥_, *z* _*f*_)d*z*_*f*_ reads,

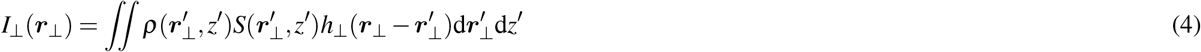

with *h*_⊥_(***r***_⊥_) = ∫ *h*(***r***_⊥_, *z*)d*z*.

We will consider two different implementations of RIM in an EDF configuration. The first one relies on the modification of the experimental system in order to generate speckles that are invariant along the optical axis. The second one uses standard three-dimensional speckles and does not require any modification of the experimental system but is only valid for a smooth surface-distributed sample.

### 2.3 EDF-RIM using speckles invariant along the optical axis

We first assume that the speckles are invariant along the optical axis, *S*(***r***_⊥_, *z*) = *S*_B_(***r***_⊥_). Such columnar speckles can be achieved experimentally by putting an annulus mask in the pupil plane to generate Bessel-type speckled illuminations^41^. In this case (see Appendix A), the theoretical expressions for the raw EDF image and its variance read,

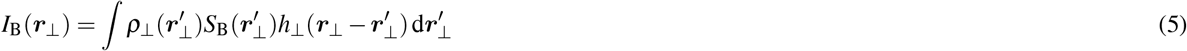

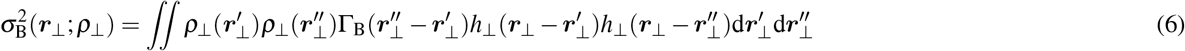

where *ρ*_⊥_(***r***_⊥_) := ∫ *ρ*(***r***_⊥_, *z*)d*z* is the projection of the fluorescence density onto the transverse plane.

In Eq. 6, we have introduced 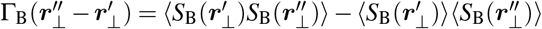 the auto-covariance of the Bessel-speckle, which is invariant along *z, i*.*e*.. In this case the EDF theoretical variance, Eq. 6, has the same structure as that derived in 2D RIM in Eq. 3. Thus we can leverage the RIM reconstruction strategy to provide a super-resolved EDF image of the projected sample *ρ*_⊥_, by changing *h* into *h*_⊥_.

Bessel-type speckles provide a clear and canonical framework to derive an EDF-RIM strategy. Obviously, whether EDF-RIM can be produced with conventional 3D-speckles is a pivotal question.

#### EDF-RIM using standard speckles for observing fluorescent surfaces

To what extent can EDF-RIM be performed with conventional 3D speckles? We recall that the EDF image depends on ∫ *ρ*(***r***_⊥_, *z*) × *S*(***r***_⊥_, *z*)dz so that the projection of the sample and that of the illumination cannot be separated. However, when the sample is a surface, (more generally a 2D manifold), EDF-RIM can be used under some approximations. We consider a sample in which the fluorophores are distributed along a surface parametrized by the function *Z*(***r***_⊥_):

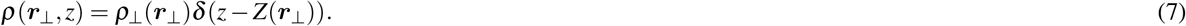

Furthermore, we assume that the fluorophores distribute on a smooth topography, implying that the surface does not vary much in *Z* over lateral distances of the order of 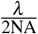 where *λ* is the wavelength of the illuminating light,

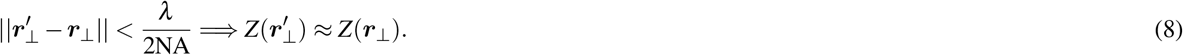

With these simplifying assumptions, we demonstrate in Appendix A that the EDF image variance also follows a 2D canonical form :

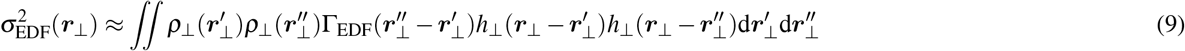

where Γ_EDF_(***r***_⊥_) is the “slice” at *z* = 0 of the 3D auto-covariance of the speckle illuminations, *i*.*e*., Γ_EDF_(***r***_⊥_) := Γ_*S*_(***r***_⊥_, *z* = 0). To conclude, we have shown that the variance of the extended depth RIM images can be simplified into the classical 2D-RIM expression of eq. 3 for any type of sample, by using columnar speckles (eq. 6), and for sparse and smoothed samples, when using regular 3D speckles (eq. 9). This result justifies that we can still use the RIM reconstruction method in this EDF configuration.

In Appendix B, we simulate EDF-RIM image formation to explore the implications of our smooth surface approximation. We simulate EDF-RIM imaging of an object with different topographies: i-a flat object for which eq. 6 is correct with no approximation; ii-a smoothly varying surface, which satisfies the conditions for the EDF variance to align with the canonical 2D-RIM expression (eq. 9); iii-a random topography, which does not satisfy the necessary conditions for the approximation. These simulations show that with a 3D propagating speckle, EDF-RIM does indeed provide super-resolution with the flat and smoothly varying topographies (conditions *i* and *ii*, see Appendix B and Fig. S1). However, with the random topography (condition *iii*), reconstruction is thwarted by amplified noise at small regularization parameter of the inversion process. Simulating EDF-RIM with a Bessel speckle, which is invariant along the optical axis, we were able to achieve super-resolved images for all samples, including the random z configuration, for every value of the regularization parameter tested (Fig. S1B, lower panels). Our simulations thus confirm that 3D-speckles can be used to image sparse samples along the optical axis, such as surfaces, and Bessel speckles can used for arbitrary samples.

In the following section, we demonstrate an experimental implementation of EDF-RIM. We place ourselves in the sparse object approximation, which allows us to use a simple 3D-speckle illumination for our extended-depth imaging. To reach an extended depth detection, we perform a focal sweep of the plane conjugated to the detector in one camera integration.

## 3 Experimental implementation

### Principle

To perform EDF-RIM in the context of the aforementioned approximation, we use fully developed speckles as 3D illuminations. The speckled illumination is then combined with a detection scheme with an extended detection PSF. This is experimentally realized by rapidly sweeping the imaged plane using an electrically tunable lens (ETL), based on a shape-changing polymer lens (see methods). A telecentric relay in the detection branch of the microscope is used to place the ETL in a plane conjugated to the pupil of the objective lens (see Fig. 1A and Methods section). We send ramps of electrical current to the ETL within a single camera exposure, effectively summing the fluorescence signal over multiple planes to achieve extended depth (Fig. 1B). With a focal power of the ETL ranging from -1.25 to 5 dpt, the focal sweep can be used for depths up to 20*µm* in ∼ 50*ms*, with our configuration.

**Figure 1.**
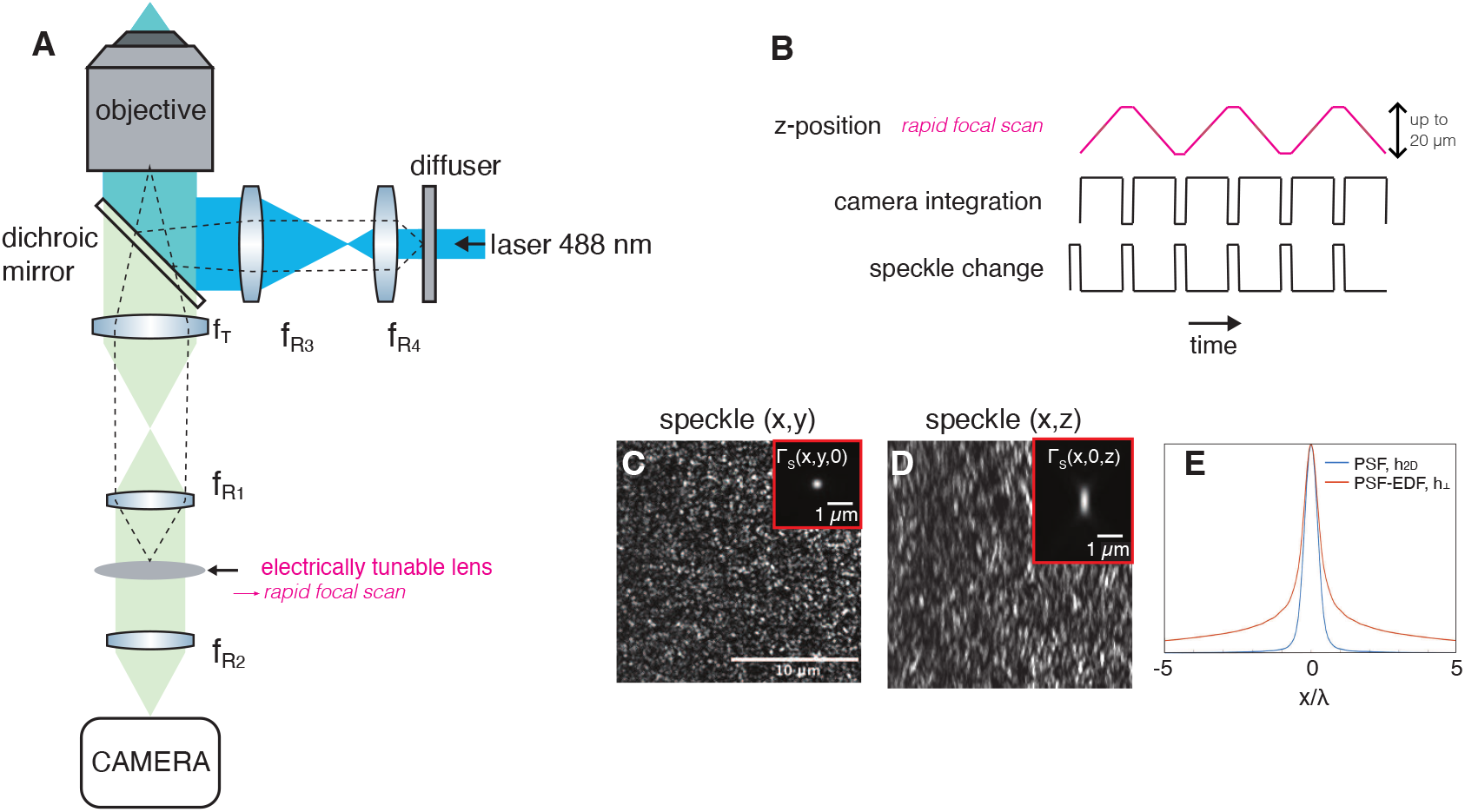
(A) Optical layout of the experimental system. Speckles are generated either with a diffuser or a spatial light modulator. The detection branch comprises a 4f optical relay (*f*_*R*1_, *f*_*R*2_) which includes an electrically tunable lens to perform a rapid focal sweep placed in a pupil plane. (B) Temporal scheme of an acquisition sequence. the top line represents the z-plane conjugated to the camera which is rapidly scanned in one integration time. (C,D) Example of an illumination speckle in the x,y and x,z planes. The insets display the speckle auto-covariance function Γ_EDF_(***r***_⊥_) := Γ_*S*_(*x, y*, 0) and Γ_*S*_(*x*, 0, *z*). (E) Comparison between the 2D PSF, *h*_2*D*_(*r*), of the wide field microscope and the EDF-PSF, *h*_⊥_(***r***_⊥_), which is obtained from the integral of the 3D-PSF along the z-axis.

Figure 1C,D show respectively (x,y) and (x,z) sections of an excitation speckle. Figure 1E shows the EDF-PSF, *h*_⊥_(***r***_⊥_). Insets in Fig. 1C,D provide respectively the auto-covariance Γ_EDF_(***r***_⊥_) := Γ_*S*_(*x, y, z* = 0) and Γ_*S*_(*x*, 0, *z*).

We determine the experimental image-variance from a set of 100-200 EDF-images captured under different speckled illuminations. The super-resolved reconstruction is then obtained through a variance matching algorithm (eq. 2), as explained in the Theory and Methods sections.

In subsequent sections, we investigate the use of EDF-RIM for super-resolved cell imaging in the context of tissue imaging.

### Application to tissue imaging

To demonstrate the effectiveness of the setup, we begin by imaging fluorescence from the actin cytoskeleton (Phalloidin-alexa488) of a human cultured cell. In Figure 2, we compare, for the same field of view and the same photon budget, the image provided by EDF-RIM (Fig. 2A), the standard EDF-widefield image (Fig. 2B) and the Wiener deconvolution of the EDF-widefield image (Fig. 2C). As expected, the raw EDF-widefield image is marred by an important background and its deconvolution efficiently improves the contrast and resolution^31^. Yet, EDF-RIM image is still significantly better contrasted and better resolved than the deconvolved widefield image, as seen in the close-up views of Fig. 2D-F and the line profiles of Fig. 2G. EDF-RIM is able to distinguish two filaments separated by 155 nm (lines 2,5) while deconvolved EDF-widefield could not distinguish the two filaments separated by 231 nm (line 3) but could distinguish the two filaments separated by 270 nm (line 4), close to the Rayleigh criterion 0.6*λ/*NA ≈ 260 nm. Thus, EDF-RIM improves the resolution by a factor close to 1.7. This resolution improvement can also be seen in the Fourier domain (Supplementary Figure S2).

**Figure 2.**
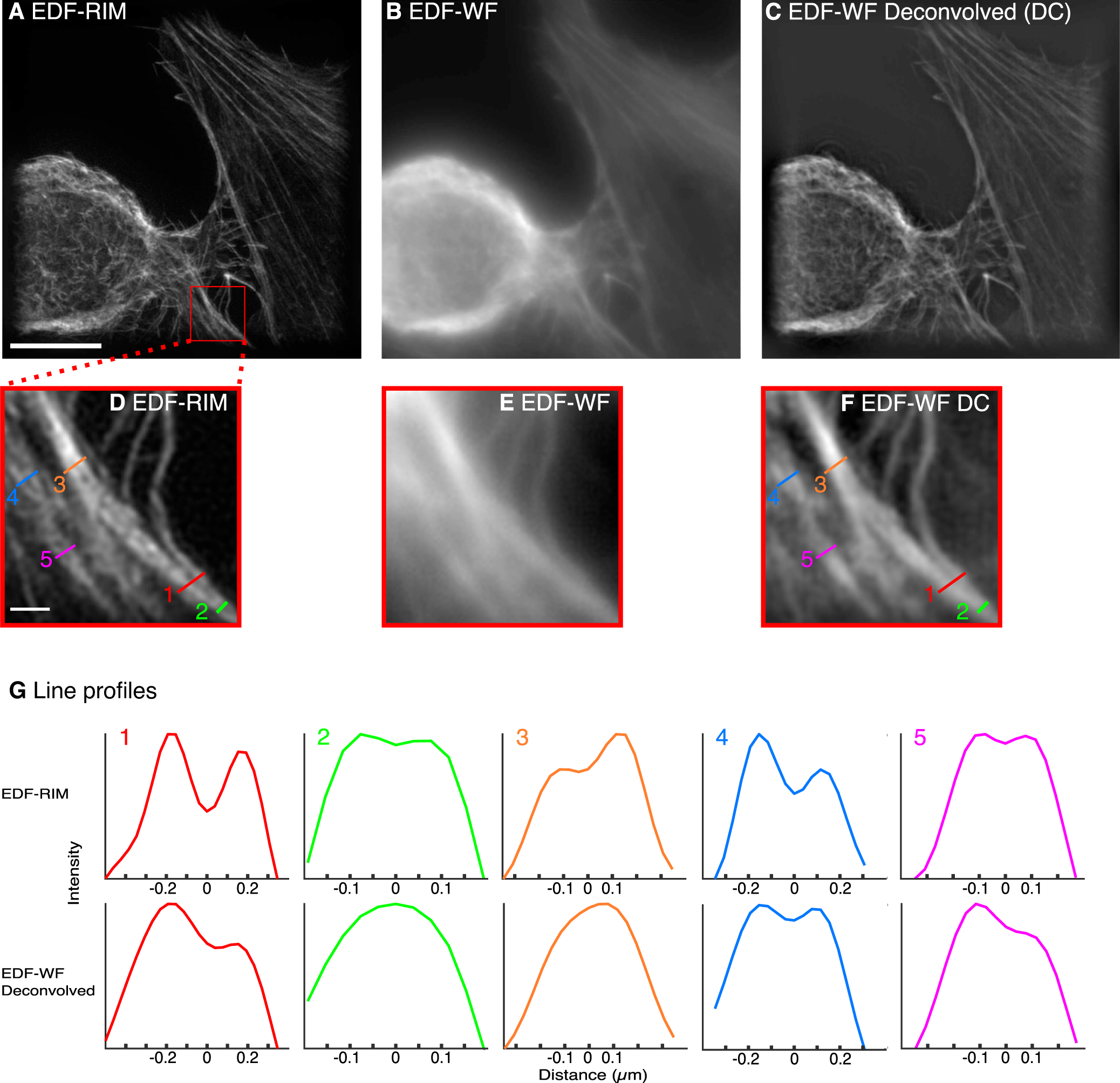
Comparison of EDF-RIM with EDF-Widefield. (A-F) Phalloidin-alexa488 labeling of the actin cytoskeleton on a cultured cell imaged with EDF-RIM (A), EDF-WF (B) and EDF-WF deconvolved (C). The projected depth is 5.5 *µ*m. Insets show a close-up view of the outlined region (D,E,F). Scale bar of the full image (A) and of the outlined region (D) are respectively 10 *µ*m and 1*µ*m. (G) Line profiles along the segments shown in EDF-RIM (D) and deconvolved EDF-WF (F). EDF-RIM is able to distinguishes two filaments separated by 155 nm (lines 2,5) whereas deconvolved EDF-WF distinguishes at best two filaments separated by 269 nm (line 4).

We further evaluate the effectiveness of EDF-RIM in tissue imaging by comparing it to 2D-RIM where different planes of the volume are acquired sequentially. We imaged desmosomes from the mouse intestinal epithelium. The structure, approximately 6*µm* deep, could be captured in a single EDF-RIM capture (Fig. 3A). By comparison, it took 14 planes to capture the same structure in conventional 2D-RIM -of which 2 planes are displayed in Fig. 3B,C. Fig. 3D displays a zoom on the EDF-RIM image. Fine details of the intestinal epithelium are visible, with a resolution comparable to the one of 2D-RIM (Fig. 3E,F). In Fig. 3D, the red and blue arrows point to structures in the single EDF-RIM image that belong to different planes, imaged sequentially in 2D-RIM (Fig. 3E,F), confirming the projective nature of EDF-RIM.

**Figure 3.**
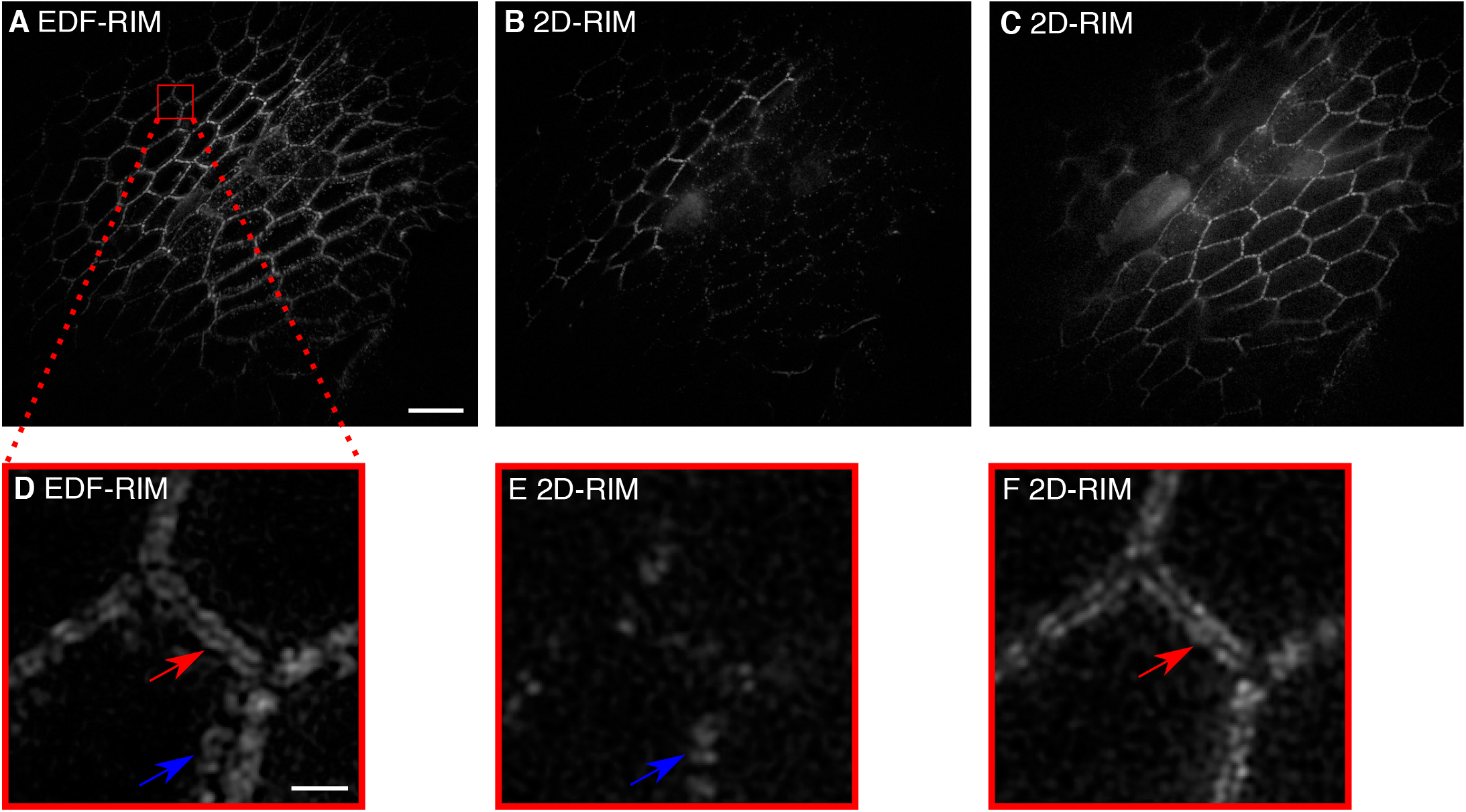
Comparison of EDF-RIM with 2D-RIM. (A) Image of desmosomes from the intestinal mouse epithelium with EDF-RIM, in which the entire 5.4 *µ*m depth is acquired simultaneously. (B,C) Two slices (z=3, and z=11) of the same tissue imaged with 2D-RIM. The volume acquisition required 14 sequential z-slices. (D) Close-up view on the EDF-RIM image. (E,F) Close-up view on 2D-RIM at planes z=2 and z=7. Red and blue arrows point to structures in the EDF-RIM image that are only visible in one of the 2D-RIM images. Scale bar of the full image (A) and of the outlined region (D) are respectively 10 *µ*m and 1*µ*m.

An essential feature of the comparison between EDF-RIM and 2D-RIM in Fig. 3A-F, is that individual images were captures with the same laser power and same camera exposure time in EDF-RIM and 2D-RIM. As a consequence, EDF-RIM implied a 14-fold reduction in light dose and acquisition time as all planes are acquired simultaneously in EDF. To further illustrate the benefit in temporal resolution, Supplementary Figure S3 and Supplementary movies 1,2 display Myosin II at the onset of mitotic domains formation in gastrulating *Drosophila* embryos. The movies are the result of a ∼ 13*µm* projection along the optical axis. This projection is performed through an EDF-acquisition in Supp. Movie 1, and by summing sequentially acquired 2D-RIM slices in Supp. Movie 2. The white arrows in both movies point to Myosin-based contractile cytokinetic rings (see the explanatory schematic in Fig. S3A). As a consequence of the increased temporal resolution in EDF-RIM, the contraction of the ring appears smooth and nicely resolved, while it appears jerky and hard to follow in 2D-RIM (Supp movie 2). The kinetic of cytokinetic ring closure displays a more than 10-fold improvement in temporal resolution (Fig S3D). In EDF imaging, information pertaining to the topography of the tissue is lost. However, it may be necessary to retain some level of topographical information, even if not at a super-resolved level. To address this, we reach a tradeoff between speed and resolution and acquire one plane-by-plane scan of the sample volume using one speckled illumination and extract topographical data with it. To achieve this, we use a robust estimation strategy, which can handle noise and background inherent to the 3D data^42^ (see Methods). The proposed approach primarily involves identifying bright points within the 3D sample and filtering out those that are not part of the most densely populated surface (Fig. 4A). Figure 4C,D shows an orthogonal view and 3D representation of the surface and associated inliers/outliers. The precision of this estimation has been investigated using information theoretical tools^43^. In Figure S4, we also show that it does not degrade rapidly as z-sampling decreases. Thus the acquisition for surface reconstruction can be fast and minimally invasive in terms of photon budget. Once the surface is estimated, we can then proceed with the regular EDF-RIM imaging protocol and ultimately project the EDF-RIM super-resolved image onto this topography for a 3D rendering of the fluorescent signal. Fig. 4E,F are respectevely the result obtained on the tight junctions from a chick neural tube and the mouse intestinal epithelium (same as in Fig. 3A-F).

**Figure 4.**
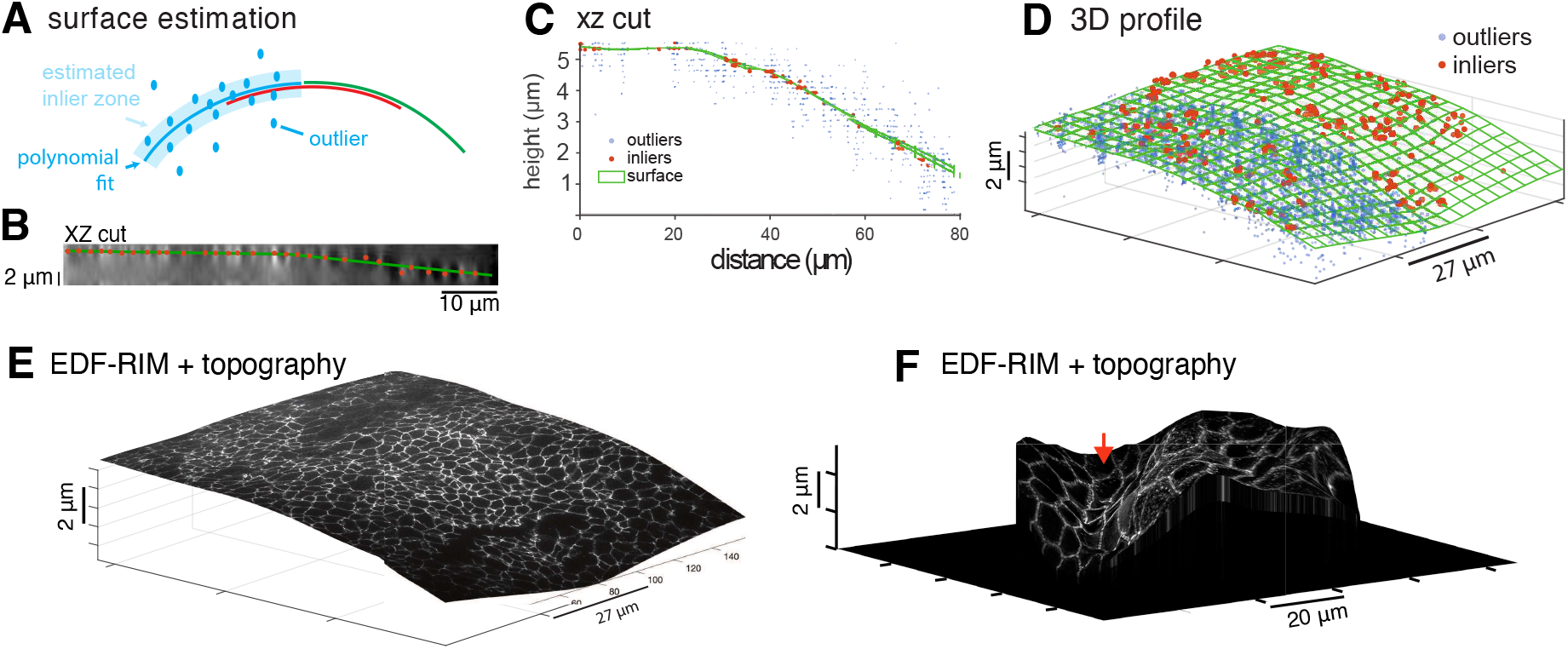
Topographical estimation. (A-E) Principle of topographical estimation illustrated on tight junctions from a chick neural tube (ZO-1 staining): one slice-by-slice stack is acquired for one speckled illumination. A consensual inlier/outlier classification of the bright points is then determined with RANSAC piece-wise fits (see methods).Inliers are then interpolated. (B) Orthogonal view of the deconvolved slice by slice stack acquired for the surface estimation (C) Orthogonal view of the surface, inliers and outliers for the chick neural tube data. (D) 3D view of surface, inliers and outliers. For sake of clarity, outliers (blue dots) are only shown on half of the surface. (E) The EDF-RIM image projected on the topography. (F) Projection of the EDF-RIM image of the desmosomes (Fig. 3A) on the estimated topography.

## 4 Discussion

In this work, we combine SIM with an extended depth-of-field approach. For this, we combined speckled illumination, extended depth of field detection and RIM treatment. Our main approach in this article is to use an ETL to perform a rapid focal sweep in one camera integration time.

The strength of EDF-RIM lies in the use of speckle as a 3D illumination function, whose statistics are invariant along the optical axis and insensitive to optical aberrations. Combined with the statistical treatment of the inversion process, this results in an insensitivity to aberrations as the system sweeps the volume of the sample —which would be difficult to achieve in a conventional SIM approach.

Classical extended-depth approaches allow an increase in acquisition speed when imaging thick samples, but they suffer from an increased background signal. We demonstrated a considerable reduction in background and an increase in resolution with EDF-RIM (Fig. 2.A,D and Fig. 3.A,D), which was also superior to a deconvolved wide-field image. Compared to 2D-RIM, EDF-RIM allows the capture of volumetric features within one acquisition. We demonstrated faster imaging of tissues with an order of magnitude gain in speed and reduction in photon budget (Fig 3. A-F, Fig S3 and Supp. Movies 1,2) Our focal sweeps were limited to ∼ 20*µm* due to the range in focal power of the ETL –still above the depths (< 13*µm*) we explored in our samples. Alternatively, deformable mirrors^31^ would also allow for faster, deeper sweeps while at the same time compensating for spherical aberrations -albeit at the expense of more costly equipment. For flat samples, another alternative is to replace the ETL with a phase mask to extend the detection PSF - thus releasing the necessity to sweep the focal plane. As an example, Fig. S5 and Supplementary movies 3,4 demonstrate EDF-RIM using a stair-step phase mask to extend the axial PSF from 0.65*µm* to 1.6*µm*. We use it to capture the rotational dynamics of Myosin-II mini-filaments in a Drosophila egg chamber. These dynamics would have been difficult to capture with an imaging system three to four times slower.

Being a projective method, EDF-RIM implies a loss of information on the biological sample under study –especially in the context of morphogenesis, where the global shape of the tissue is an essential feature. In these circumstances, the low-resolution topographical estimation that we propose (Fig. 4E-F), which requires one slice-by-slice acquisition, may represent a satisfactory compromise between speed of acquisition and proper description of the geometrical features of the biological sample.

What are the limits of our current approach? We made the assumption that the fluorophore density is sparse along the optical axis. This allows us to reach a simple model for image formation that we use in the inversion process (see Theory section). A typically good configuration is when fluorophores are distributed over a smoothly varying surface (or several surfaces). Conversely, a randomly distributed topography can lead to a greater sensitivity to noise of the reconstruction at low regularization parameter, as demonstrated in our simulations (Fig. S1B, top-right panels). Experiments, however, tell us that the restriction is not drastic. We could image at high resolution the actin cytoskeleton of an entire cell, which is ∼ 5*µm* thick, with a much-improved contrast and resolution over extended wide field (Fig. 2A-F).

EDF-RIM also suffers from the same limitations as RIM and SIM methods in general. A primary source of reconstruction error inherent to all SIM approaches stems from photon noise originating from areas outside the plane of focus (or outside the integrated volume in the context of EDF). The magnitude of this noise from the out-of-plane blur, relative to the fluorescence fluctuations tied to the structured illumination (whether harmonic-based or speckle-based), directly impacts the reconstruction error. Another source of noise common to all RIM techniques comes from the discrepancy between the empirical variance (the variance estimated from a limited number of speckles, typically 100 200 in our experiments) and the asymptotic variance we would obtain with an infinite number of speckles. This introduces a residual granularity that can affect the final resolution. Finally, RIM techniques don’t demand more photons compared to other SIM methods or even widefield imaging. Our comparisons with widefield were conducted under a constant photon budget (Fig. 2A-C). Consequently, the photon count per image in RIM is lower, and potentially susceptible to camera read noise in low photon flux scenarios. Thus, EDF-RIM, which reduces the number of images required via projective imaging, may present an enhancement in this regard.

One improvement in future implementations could be the use of an illumination that is truly invariant in z, not just statistically invariant. This could be done using Bessel speckles –achieved experimentally by blocking the light outside an annulus in the pupil plane, and maintaining a random phase within this annulus with a diffuser or spatial light modulator. This would improve the inversion process for samples that are not distributed over a surface (Fig. S1B, bottom-right panels). The approach could also improve resolution owing to the stronger content in high spatial frequencies of Bessel speckles^41^.

Projective approaches have been recently combined with SIM in light sheet microscopy^44^. Here, we have developed the first wide-field structured illumination microscope, which combines super-resolution with an extended depth capacity. For this, we build upon the framework of the random illumination microscope (RIM), combining the intrinsically 3D nature of illumination speckles with extended depth detection. EDF-RIM appears as an attractive alternative to the classical EDF-widefield when a high resolution and low background is necessary, for example in the context of high-resolution molecular tracking and tissue imaging.

## Methods

### Optical setup

The two approaches have been built on separate setups for sake of convenience, but share a similar optical layout. Essentially, the intermediate image created by the objective and tube lenses is relayed by *f*_*R*1_ and *f*_*R*2_, arranged in a 4 *f* configuration, onto the camera. An objective and a tube lens create a magnified image of the sample (*M* = *f*_*T*_ */ f*_*O*_) in an intermediate image plane. The intermediate image is relayed by *f*_*R*1_ and *f*_*R*2_, arranged in a 4 *f* configuration, onto the camera. The ETL or phase mask are placed in a pupil plane in between *f*_*R*1_ and *f*_*R*2_^45,46^. Excitation speckles are characterized (auto-covariance function Γ_*S*_(*x, y, z*)) by replacing the sample with a 50 nm gold film and using the remote focus to image the speckle in 3D.

#### Rapid focal sweep set-up

The microscope is custom built, made of optomechanical elements. *f*_*O*_ is an 100x objectify (Nikon, N100X-PFO, nominal NA=1.3, effective NA=1.21 for Fig. 2A-F; CFI SR APO 100× NA 1.49 Nikon for Fig. 3A-F), *f*_*T*_, *f*_*R*1_, *f*_*R*2_ are thorlabs TTL165-A, AC254-125-A-ML,AC254-125-A-ML. The exact position of the ETL and its associated offset lens (Optotune EL-10-30-Ci Series) is finely adjusted so as to optimize the detection PSF. We found that placing specifically the ETL in the fourier plane and not the offset lens or the mid-point between the offset lens and the ETL was optimal. The lenses *f*_*O*_, *f*_*T*_, *f*_*R*1_, *f*_*R*2_ are arranged to ensure telecentry of the imaging path. In the excitation branch of the set-up, light from a 488 nm diode laser (Oxxius, LBX-488-200-CSB), impinging on a diffuser or a SLM, is relayed to the pupil of the objective by *f*_*R*3_ and *f*_*R*4_ (Thorlabs LA1986-A-ML and LA1131-A-ML). With this configuration, the maximal range of the ETL focal power (−1.5 to 4.5 dpt) lead to a maximum z-range of 21*µm* in the sample space. Successive speckles are triggered through rotation of the diffuser with a rotary stage (2-axis controller 8SMC4-USB-B9-2) in Figs 2A-F and with a spatial light phase binary modulator (SLM, QXGA Fourth Dimension) for Fig. 3A-F. The illumination beam is combined with the imaging path with a dichroic miror (semrock, Di02-R488-25×36). Fluorescence is further filtered with a bandpass filter (Semrock, FF01-525/45-25) and imaged with a camera (Prime BSI Express Scientific CMOS Camera). A Matlab interface controls all instruments of the setup. Hardware synchronization of instruments is done using a National Instrument (NI USB-6351) board, though digital pulses and analog signal sent to the ETL driver.

#### Phase mask setup (Fig. S5

An inverted microscope (TEi Nikon) is used with a 100× TIRF objective (CFI SR APO 100× NA 1.49 Nikon) We use a 488 nm diode laser (Oxxius, LBX-488-200-CSB) for GFP/Alexa-488 and a 561 nm solid-state laser (Oxxius, LMX-561L-200-COL) for cherry imaging. A fast spatial light phase binary modulator (SLM, QXGA Fourth Dimension) placed in an image plane was illuminated by the collimated 8.8 mm TEM00 beam. Fluorescence, after the dichroic mirror (Semrock DI02-R488-25×3625×36) and Stop-Line notch filter (Semrock NF03-405/488/561/635E-25) is collected by an sCMOS camera (OrcaFlash fusion). The stair-step phase mask (DoubleHelix, PNDT6) is placed in a pupil plane of the optical really consisting of *f*_*R*1_, *f*_*R*2_ (focal lengths: respectively 200 mm and 300mm). Two bandpasses filter the collected fluorescence (Semrock FF01-514/30-25 for green fluorescent protein and Semrock FF01-650/92 for cherry). User-interface and synchronization of all instruments are provided by the INSCOPER software and controller.

### Image reconstruction

The reconstruction is essentially based on the algoRIM treatment^47,48^ developed for the analysis of 2D-RIM images, and which is accessible at https://github.com/teamRIM/tutoRIM. In brief, the N images taken under different speckled illumination (*N* ∼ 100 − 200 in most cases) are first processed with a Wiener filter, which can be written in the Fourier domain:

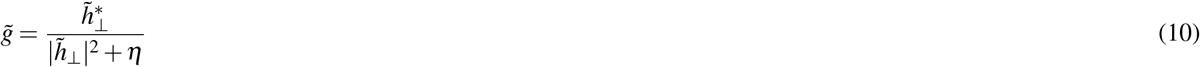

where 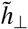 is the Fourier transform of the extended depth PSF, *a*^*^ stands for the complex conjugate of *a*, and *η* is a regularization parameter set at 10^−3^. This filter enhances high frequencies inside the OTF and removes noise beyond the OTF cutoff. We then estimate empirically the standard deviation of these pre-filtered speckled images. Finally, an iterative, moment-matching algorithm is used to achieve a super-resolved reconstruction of the object. We use a Tikhonov regularization method such that the reconstructed fluorescence density is given by

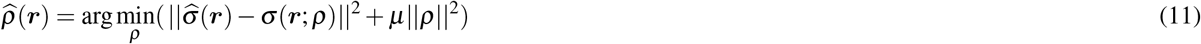

with 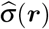 the experimental standard-deviation and *σ* (***r***; *ρ*) the theoretical standard-deviation derived from the expected variance. In this theoretical expression, we generate a auto-covariance function of the speckle and PSF that matches the Fourier support of the experimental auto-covariance function of the speckle. The difference with eq. 2 stands in the presence of the regularization term *µ*||*ρ*||^2^ in which *µ* is the Tikhonov regularization parameter. Due to the preprocessing step, we modified the model of the theoretical variance (eq. 6 for columnar speckles, eq. 9 for 3D speckles) by replacing *h*_⊥_ with *h*_⊥_ * *g*. When set too high, the regularization parameter *µ* can lead to image blurring and prevent super-resolution. Conversely, when set too low, it may lead to the amplification of noise and the appearance of artifacts. For experiments, we used a value *µ* = 10^−5^ (Fig. 2A). In the simulations (Appendix B), we explore the link between the value of *µ* and performance of the reconstruction in situations where the model for EDF image formation is correct, and when it is not. For the cell data (Fig.2 A-F) we applied to the raw data a Tukey window with cosine fraction r=0.2 to reduce the Gibbs phenomenon. In Fig.3 A-F, we remove a smooth, out of volume of interrogation, background with a rolling ball algorithm^49^. For the deconvolution of the EDF-WF image, we used a Wiener filter (10). The regularization parameter (*η*) was set at 2.10^−4^ by eye to get the same level of artefacts than the EDF-RIM for a fair comparison.

### Estimation of surface topography from one full scan

To estimate surface topography, the remote focus is used to capture the fluorescence signal under one speckled illumination plane by plane (*ie* one image per plane). This is typically performed by acquiring one plane every 2 *µm*. We then employ a robust approach for estimating surface topography, based on the method presented by Abouakil et al.^42^ and Meng et al.^50^. To correct for the spatial inhomogeneities of the background signal, we introduce a normalized signal of the form:

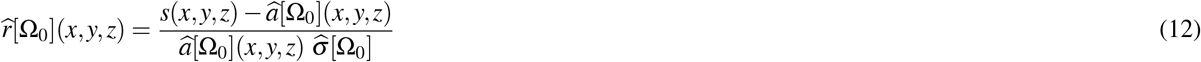

Here, *s*(*x, y, z*) is the measured signal, 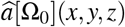 is an estimation of the spatial inhomogeneities of the background, and 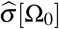 is an estimation of the standard deviation of 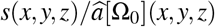 on the background. We determine both 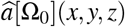 and 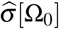 using only a subvolume Ω_0_ of the acquired volume, taking randomly a small fraction (1/1000) of the acquired voxels. This treatment allows for a faster surface estimation.

The resulting normalized signal has a heavy-tailed Gaussian-like distribution in the histogram, with the Gaussian representing the background and the heavy tail representing the signal of interest from the biological structure. We set a threshold based on a chosen probability of false alarm (*p* = 0.01) and interpolate the epithelial surface modeled as *z* = *Z*_*s*_(*x, y*) using the detected bright points. To estimate the surface over large regions of interest, we use quadratic polynomial fits in overlapping windows (typically 3×3 to 5×5 windows), then use the RANSAC technique to estimate the most consensual classification of inliers and outliers^51^. We fuse the overlapping estimations by preserving the inliers in all of the overlapping quadratic fits while discarding the remaining inliers. Finally, we estimate the surface by interpolating the remaining inliers using a simple bi-cubic harmonic spline interpolation.

### Simulations

For the simulations, the object was defined with a 512x512x128 grid, with *d*_*x*_ = *d*_*y*_ = 38.5*nm* and *d*_*z*_ = 87.5*nm* that ensured Nyquist criterium on the super-resolved simulated reconstruction. The wavelength in water was 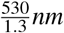 and the numerical aperture was 0.8. The simulated object is the combination of a star-shaped lateral distribution *ρ*(*r, θ*), and a topography *Z*(*r, θ*) (Fig. S1A). The lateral distribution follow the equation *ρ*(*r, θ*) = 1 + *cos*(40*θ*), which generates a star pattern characterized by higher spatial frequencies towards the center of the patter. The topography follows three different shapes: i) a flat constant *Z*; ii) a dome-shaped topography corresponding to a right circular cone with its radius equals to 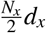, its height equals to *N*_*z*_*d*_*z*_ for 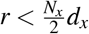 and a flat constant *Z* for 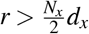; iii) a random topography with a uniformly distributed probability over the entire z-range, and no correlation between neighboring (*x, y*) points.

To generate illumination speckles, we computed the propagated field with a mask in the pupil plane with random phase and binary amplitude in the form of a disc for 3D-speckles and an annulus for Bessel speckles. The propagated speckle is given a Gaussian envelope, which introduces a correlation length in the spatial distribution of the random phases. A 3D image is then formed by multiplying the squared amplitude of the field with the object and convolving with the 3D PSF corresponding to the numerical aperture of the pupil. The EDF image is generated by summing the 3D image over all z. The variance was estimated using 1000 prefiltered images. Simulations were run in Matlab.

### Samples preparation

#### Immunolabeling of mouse intestinal epithelium

The jejunum of a sacrificed mouse was fixed in 4% paraformaldehyde (Electron Microscopy Sciences), washed in PBS, cut in 200-300 µM thick slices, permeabilized with %1 Triton X-100 for 1h. Desmoplakin immunolabeling was done through overnight primary antibody incubation (Progen, 651109, 1/50 dilution), followed by washes in PBS, overnight secondary antibody incubation (Goat anti-Mouse, Alexa Fluor-488, Invitrogen, A11029, 1/200 dilution), washes in PBS and mounting in Aqua-Poly/Mount, (Polysciences).

#### Drosophila egg chamber live imaging

We followed the protocol of Prasad *et al*.^52^. Briefly, Drosophila egg chambers (stage 9 to stage 10A) were dissected from adult females and mounted in Schneider’s insect medium supplemented with 20% FBS and adjusted to a pH of 6.9. Prior to imaging, the egg chambers were slightly flattened on the coverslip by gentle pressure. Myosin-II was imaged in a transgenic line that expresses a protein fusion of Cherry with the regulatory light chain^53^ (*squash* gene) sqh:Cherry

#### Cytoskeletal labeling of cultured cells

cells were fixed with 4% Paraformaldehyde HCHO (15714 Electron Microscopy Sciences) in 37°C-prewarmed cytoskeleton buffer (10 mM MES with pH 6.1 with NaOH, 150 mM NaCl, 5 mM EGTA, 5mM glucose, 5mM *MgCl*_2_) for 15 minutes and rinsed with PBS. Cells were then permeabilized, exposed to blocking solution, incubated with primary and secondary antibodies together with 0.165*µM* Alexa Fluor 488-phalloidin (A12379 Thermo Fisher Scientific) for 2 h at RT. While cells were treated with primary and secondary immunolabels, in this work we only imaged 488-phalloidin for actin cytoskeleton labeling.

#### Tight junction staining of chick neural tube

Chick embryos were fixed for 1 h in ice-cold 4% formaldehyde in PBS. Then they were cut along their midline and permeabilized for 15 min in 0.3% Triton X-100 in PBS (PBT 0.3%), before a 1-h blocking step in PBT 0.3% with 10% Goat Serum. The primary antibody is a mouse anti-ZO1 (Invitrogen ZO1-1A12) at 1:250 dilution, and secondary antibody is a goat anti-Mouse-alexaFluor488 (ThermoFisher Scientific, A28180) used at 1:500 dilutions. Embryos were then mounted between slide and coverslip using Vectashield (Vector Laboratories).

## Acknowledgments

We thank Sophie Brasselet, Sandro Heuke and Hervé Rigneault, for support and fruitful discussions on the project. Fig. S3A was reated with BioRender.com

This work was funded by the following agencies: Agence Nationale de la Recherche (ANR-18-CE13-028, ANR-20-CE45-0024, ANR-22-CE13-0039, ANR-22-CE42-0010, ANR-22-CE42-0026); Institut Carnot star (3D-RIM).

This project is funded by the ≪ France 2030 ≫ investment plan managed by the French National Research Agency (ANR-16-CONV-0001, ANR-21-ESRE-0002), and from Excellence Initiative of Aix-Marseille University - A*MIDEX.

## Supplementary Material

Appendix A: Detailed theory of EDF-RIM

Appendix B: Simulations

Supplementary Figure S1: Simulations

Supplementary Figure S2: Fourier spectra analysis

Supplementary Figure S3: Cytokinesis quantification in the *Drosophila* embryo

Supplementary Figure S4: Effect of z-sampling on the quality of surface reconstruction

Supplementary Figure S5: EDF-RIM using a stair-step phase mask

Supplementary Movie caption SM1

Supplementary Movie caption SM2

Supplementary Movie caption SM3

Supplementary Movie caption SM4

## Appendix A: Detailed theory of EDF-RIM

### Model for extend depth image formation

We write ***r*** := (***r***_⊥_, *z*) ∈ ℝ^3^ as a spatial coordinate in 3D, with ***r***_⊥_ := (*x, y*) locating a position in the (transverse) plane perpendicular to the optical axis *z*. Additionally, if *f* is a function defined over of ***r***, we note *f*_⊥_ its projection along the optical axis

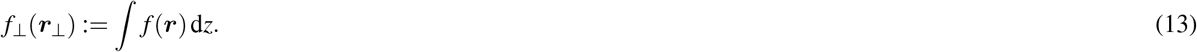

For a given 3D speckle excitation, the EDF (2D) image corresponds to the sum of the contribution from all planes, which writes as follows:

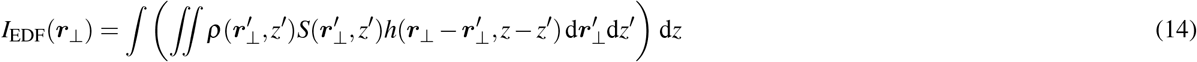

with *h* is the 3D PSF of the microscope, *S* the speckled illumination and *ρ* the fluorescence density (which is the product between the fluorophore concentration and their brightness). In the above equation, if we integrate along *z* first, we get

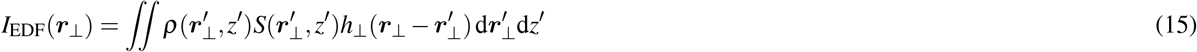

where we used the fact that *h*(***r***_⊥_, *z* − *z*^′^)d*z* = ∫ *h*(***r***_⊥_, *z*)d*z* = *h*_⊥_(***r***_⊥_). Let us assume further that speckles are columnar functions. In such an instance, *S* is invariant along *z* and we write *S*(***r***_⊥_, *z*) = *S*_B_(***r***_⊥_), ∀*z*. The relation above has the following specific form

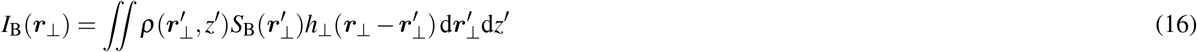

and if we finally integrate along *z*^′^, we obtain a simple convolution model that involves projected quantities only

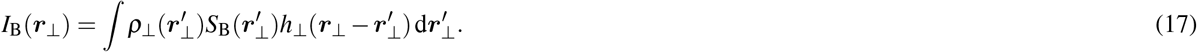

In summary, under the assumption of a Bessel illumination, the model for the microscope image is analogous to the one of a standard widefield microscope, but it involves the extended depth PSF, *h*_⊥_ and the projection of the object *ρ*_⊥_.

### Theoretical expression of the EDF variance

We derive now the theoretical variance of a EDF-RIM experiment. We start first by making no specific assumption about the illumination function *S*(***r***_⊥_, *z*). More specifically, we write from Eq.15:

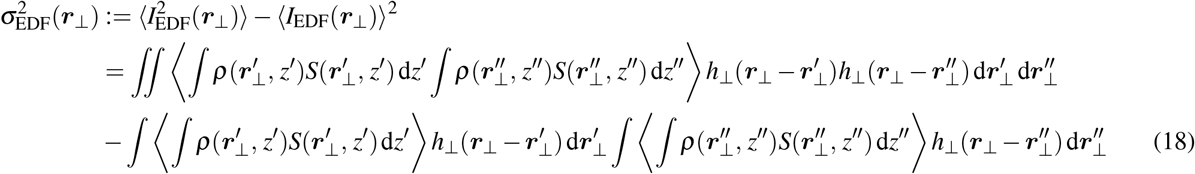

Assuming speckles that are invariant along *z*, we can write *S*(***r***_⊥_, *z*) = *S*_B_(***r***_⊥_) which allows the following simplification

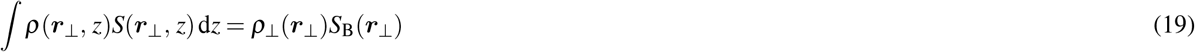

hence leading to the following approximation for the EDF-RIM variance

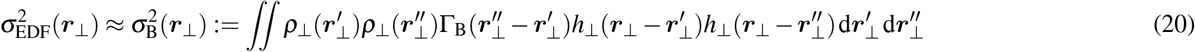

where we have introduced the 2D auto-covariance of the columnar speckle 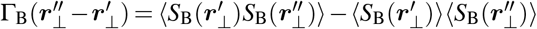. The right-hand side in Eq. 20 has the structure of the variance expression in RIM given by Eq. 3. It should be noted, however, that the classical result *h*_⊥_ = Γ_B_(***r***_⊥_) will not be verified, unless we modify the observation PSF by putting a ring in the pupil (Fourier plane).

### EDF-RIM variance with standard speckles: the case of a spatially smooth fluorescence surface

Bessel-speckles are rarely used as an illumination function. Here, we investigate under what conditions EDF-RIM can be performed with conventional 3D speckles. We thus return to the general case of an illumination speckle that is not invariant along *z*. Using Eq. 18, the EDF-RIM variance reads

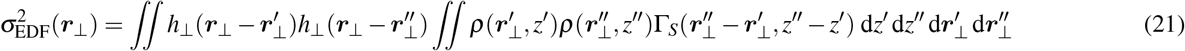

where we have introduced the 3D auto-covariance of the speckle 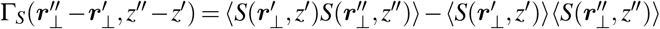. Let us assume now that the sample is such that the fluorophores are distributed along a surface denoted *Z*(***r***_⊥_):

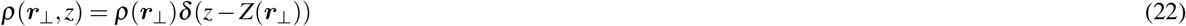

where *δ* is the Dirac distribution. Under this assumption, Eq. 21 simplifies into

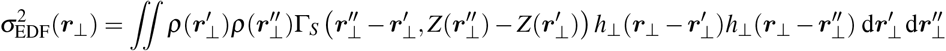

This expression is similar to the standard 2D RIM expression of the variance Eq. 9 except that it involves 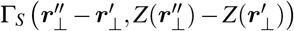. rather than 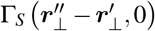 To further simplify the speckle auto-covariance into a 2D expression, we hypothesize that the objects distribute on a smooth topography. This translates mathematically into the following assumption for Γ_*S*_:

- 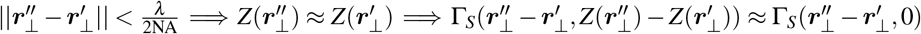
- 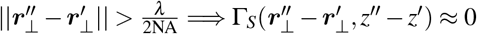

Two assumptions have been made. The first one states that the surface does not vary much in *z* over lateral distances of the order of 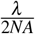 (smoothness hypothesis). The second one neglects long-range correlations in the illumination, which is ensured because speckle decorrelates on length-scales similar to the lateral extent of the PSF, 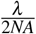^54^. Following these assumptions, the expression of the variance of the intensity is simplified: it involves the 2D PSF *h*_⊥_ and the 2D auto-covariance function Γ_EDF_(***r***_⊥_) := Γ_*S*_(***r***_⊥_, 0):

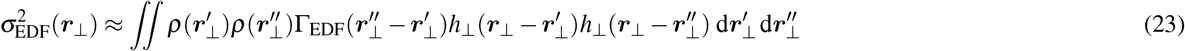

To conclude, our study has shown that the variance of the extended depth RIM images can be simplified into the classical RIM 2D expression of Eq. 3 for any type of sample, by using columnar speckles (see Eq. 20), and for surface and smoothed samples, when using regular 3D speckles (see Eq. 23).

In the following section, we demonstrate an experimental implementation of EDF-RIM. We then investigate the performance of EDF-RIM and the limits of its numerical approach through simulations.

## Appendix B: Simulations

We investigate the realms of application of EDF-RIM through simulations. More specifically, we examine the implications of the EDF variance expression in the case of the 3D speckle (eq. 21) only approximately conforming to the canonical 2D-RIM expression (eq. 3). In the simulations, the imaged object is characterized by its star-shaped lateral distribution *ρ*(*r*), as shown in Fig. S1.A (left). We investigate three distinct topographies for the 3D object, as depicted in Fig. S1.A (right). These include a flat structure, a smoothly varying surface in shape of a cone, and a random distribution of z-position. The flat object serves as a reference for which our model is correct, with no approximation. The smoothly varying surface satisfies the approximation conditions for the EDF variance to align with the canonical 2D-RIM expression (eq 9), whereas the random configuration does not.

From a numerical standpoint, the inversion requires the deconvolution of the variance of the prefiltered images (*η* = 10^−6^) images using a Tikhonov regularized inverse filter^27^. The effectiveness of this inverse filter depends on its regularization parameter. When this parameter is too high, it can result in image blurring and prevent super-resolution. Conversely, setting it too low may lead to the amplification of noise and the appearance of artefacts. We compared the inversions for 2 different regularization parameters -one large (10^−5^) and one low (10^−10^ respectively), and for the 3 aforementioned 3D objects, the idea being that an inaccurate model generally induces more artifacts for small regularization parameters.

We first examine the scenario of a 3D propagating speckle, which aligns with our experimental implementation. In Fig. S1.B, we present the reconstructed images for the three topographies, with both large (*µ* = 10^−5^) and small (*µ* = 10^−10^) Tikhonov regularization parameters. While all three results appear satisfactory at *µ* = 10^−5^, this setting is over-regularized, leading to a suboptimal resolution. At *µ* = 10^−10^, only the smooth surface compares well with the 2D case, while reconstruction of the random z-position is thwarted by amplified noise. Thus, it stands that conditions that permit the use of an exact or at least approximate expression for the image variance are a prerequisite for EDF-RIM.

As outlined in Appendix A, the EDF-variance expression can be transformed into a 2D-RIM expression for all types of samples when using a speckle that remains invariant along the optical axis. This condition is achieved experimentally by blocking light outside an annulus in the pupil plane (Bessel speckle, Fig. S1.C, right). We simulated these Bessel speckles by modifying the binary mask in the pupil plane which is no longer a disk but an annulus. This annulus filters out all frequencies smaller than 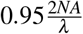 and bigger than 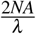. The generated speckle are effectively invariant along the optical axis (Fig. S1.C, right panels).

Simulating EDF-RIM with these Bessel speckles, we were able to achieve high-resolution images for all samples, including the random z configuration, for every value of the regularization parameter tested (Fig. S1.B, lower panels). This is consistent with our prediction that with columnar speckles, the reconstruction does not rely on the sparsity hypothesis.

In conclusion, our simulations confirm that EDF-RIM is a feasible imaging method, and support the limits outlined in the theory section. Specifically, we have demonstrated that 3D-speckles can be used to image sparse samples along the optical axis, such as surfaces, and Bessel speckles can used for arbitrary samples.

**Figure S1.**
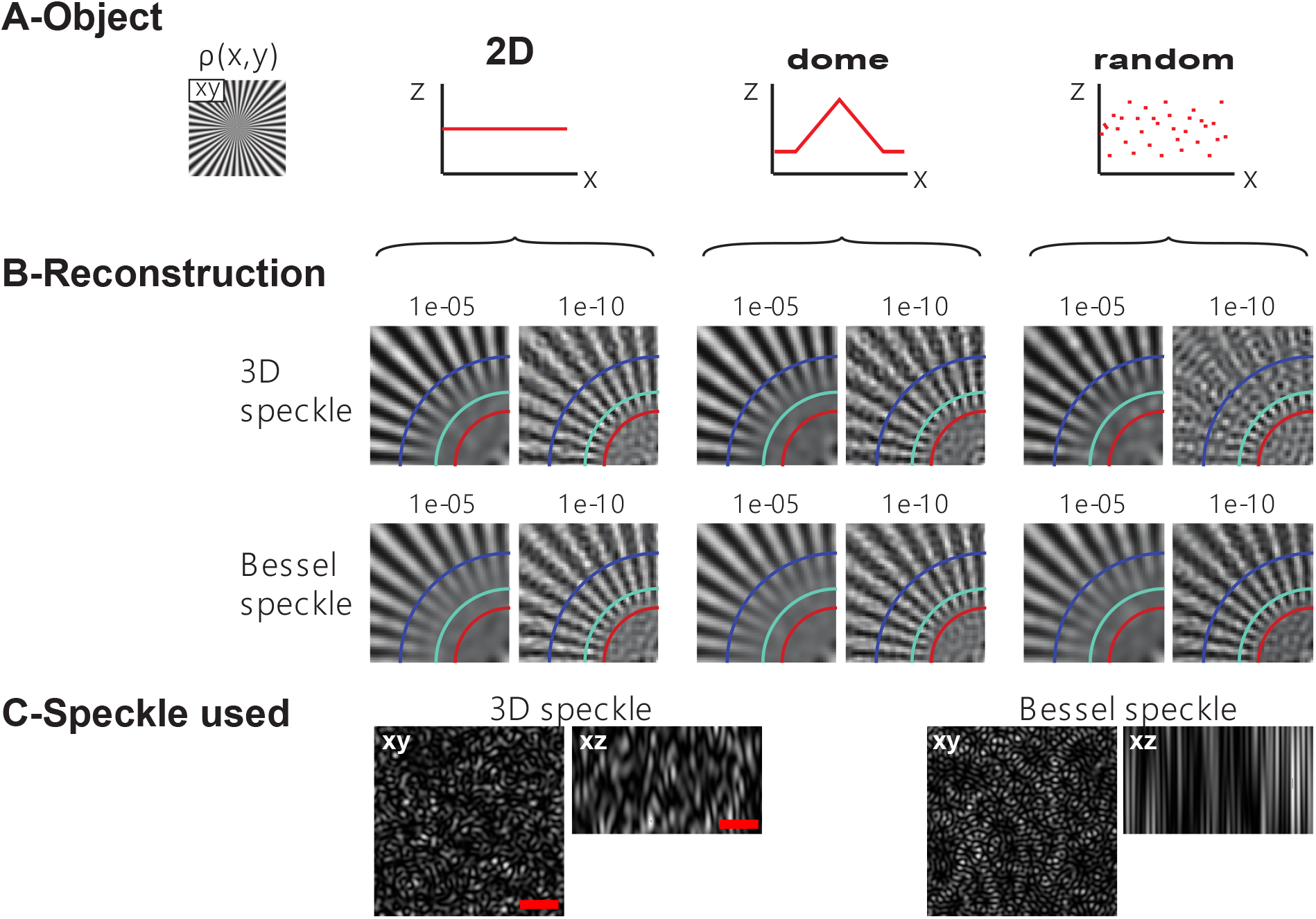
Simulations. A) Simulated object: the lateral distribution *ρ*(*r*) is depicted on the left, while three topographies are studied – a flat structure (left), a dome (center) and a random topography (right). B) Simulated reconstructions for the 3 topographies studied, with 2 different illumination-patterns (see below). For each condition, we consider the reconstruction using a large and a small regularization parameter in the inversion procedure (10^−5^ and 10^−10^ respectively). Images include guidelines at 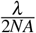 (blue circle), 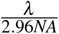 (cyan circle) and 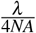 (red circle). C) The three dimensional illumination patterns used in the simulation (scale bar: 2 *µ*m). The 3D-Speckle is obtained numerically by simulating the propagation of an electromagnetic field with uniform amplitude and random phase distribution in the back focal plane of the objective. The Bessel speckle is obtained by replacing the homogeneous amplitude in the back focal plane of the lens with an annulus illumination.

**Figure S2.**
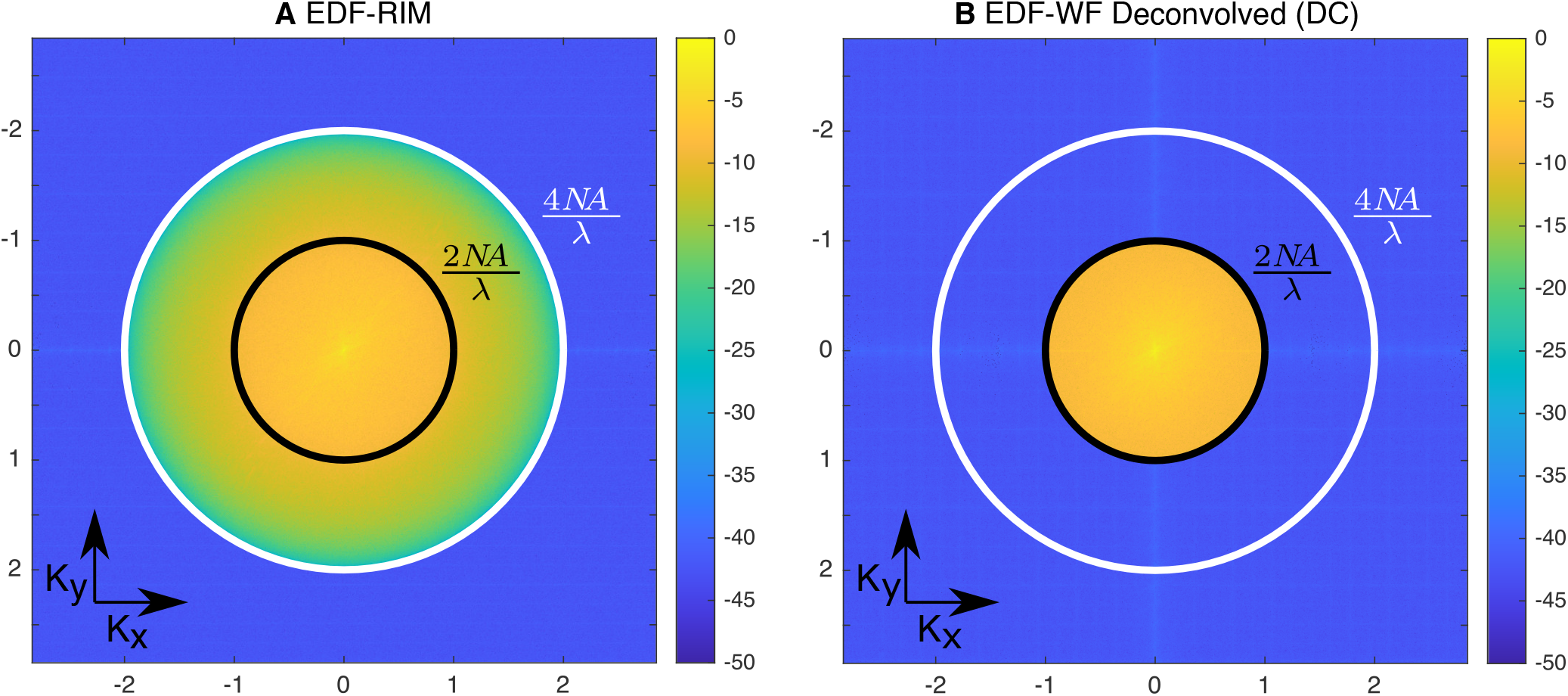
Fourier spectral analysis. (A, B) correspond to the Fourier spectra in log scale of Fig. 2.A,C. Spatial frequencies are normalized by the OTF cut-off frequency 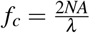 with *NA* = 1.21 and *λ* = 530 nm. The black and white circles indicate the frequencies *f*_*c*_ and 2 *f*_*c*_, respectively. (C) Line profile along *K_x_* for *K_y_* = 0 of the EDF-RIM Fourier spectra (blue curve) and deconvolved EDF-widefield (red curve). The Fourier spectrum of deconvolved EDF-widefield is limited to *f*_*c*_ while the Fourier spectrum of EDF-RIM is limited to 2 *f*_*c*_. There is an intensity decay in the Fourier spectrum of EDF-RIM at roughly 1.7 *f*_*c*_. By using smaller regularization parameters, this value could be improved to be closer to 2 *f*_*c*_, but would generate more artefacts.

**Figure S3.**
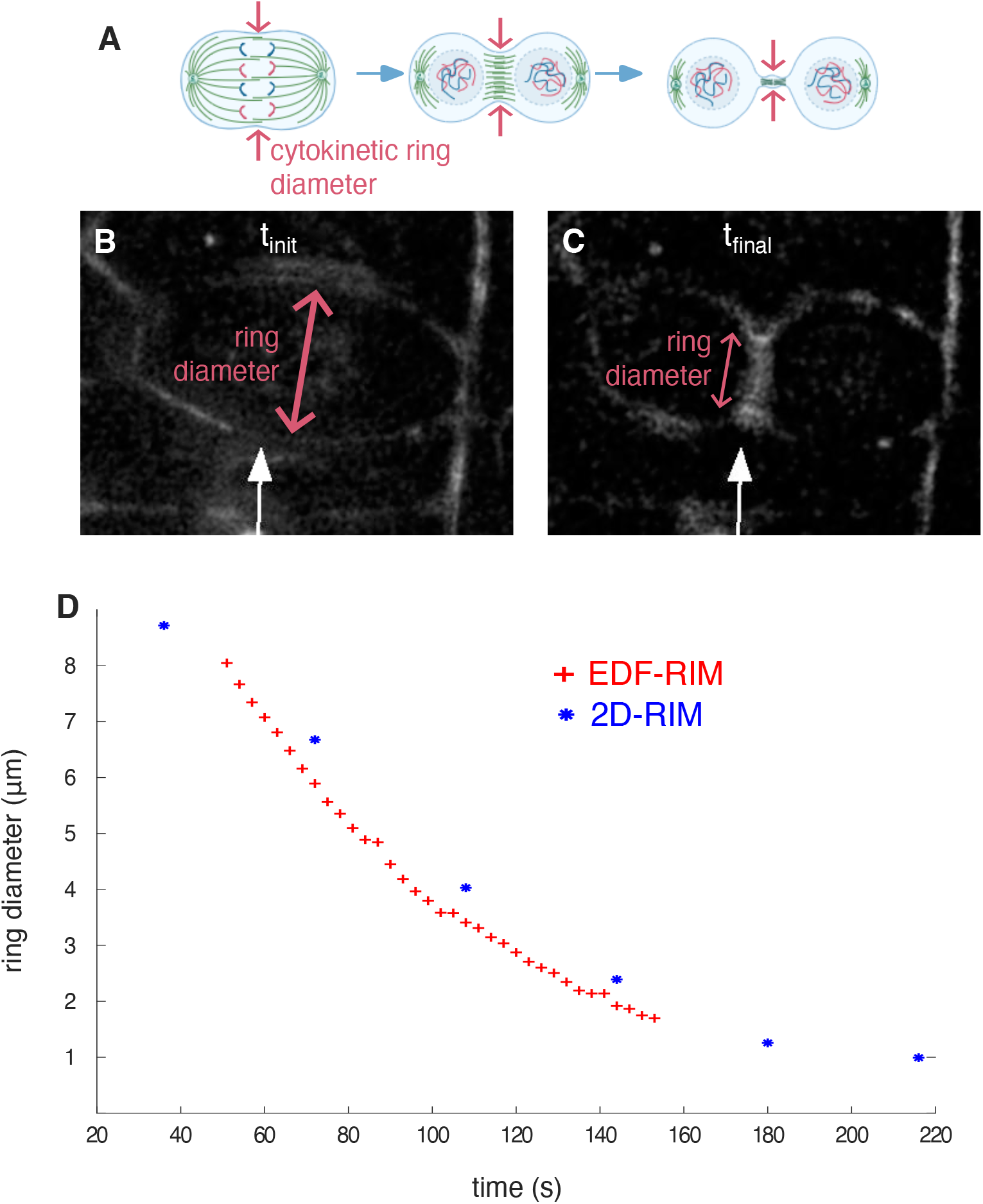
Cytokinesis quantification in the *Drosophila* embryo. Quantification of cytokinetic ring contraction taken from Supplementary movies 1. A) Schematic of a dividing cell showing the cytokinetic ring, measured in this figure. B,C) Snapshots from Supplementary movie 1, in which the white arrows point to dividing cells. D) Ring contraction as measured from Supplementary movie 1 using EDF-RIM and Supplementary movie 2 using 2D-RIM. Each point corresponds to one time point from the movie. The superior temporal resolution of EDF-RIM is reflected in the more than 10-fold increased density of measurement points. The reduced temporal resolution in 2D-RIM results in jitter, than makes cell divisions hard to follow (white arrow in Supplementary movie 2).

**Figure S4.**
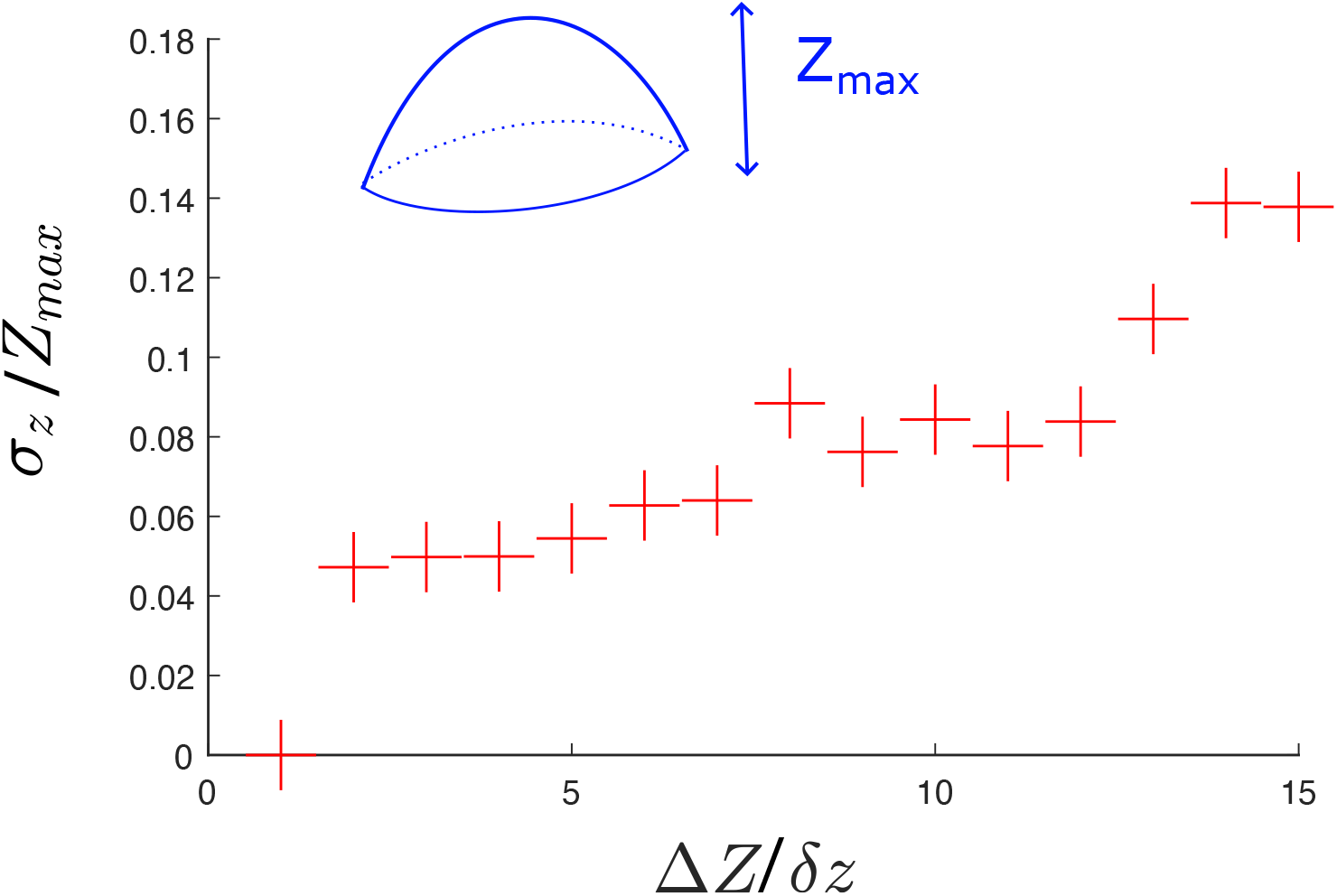
Effect of z-sampling on the quality of surface reconstruction. To estimate the dependence of the surface reconstruction on z-sampling, we estimate the surface using increasing Z intervals Δ*Z* = *n*.*δz*, where *δ* = 177*nm* is the minimal interval used, and n is an integer. The error with respect to the best estimation (n=1), is quantified as the root mean squared normalized by the height of the tissue (*Z*_*max*_, see inset on top left of image). The z-dependence is quite low up to *n* = 12 corresponding to Δ*Z* ≃ 2.1*µm*.

**Figure S5.**
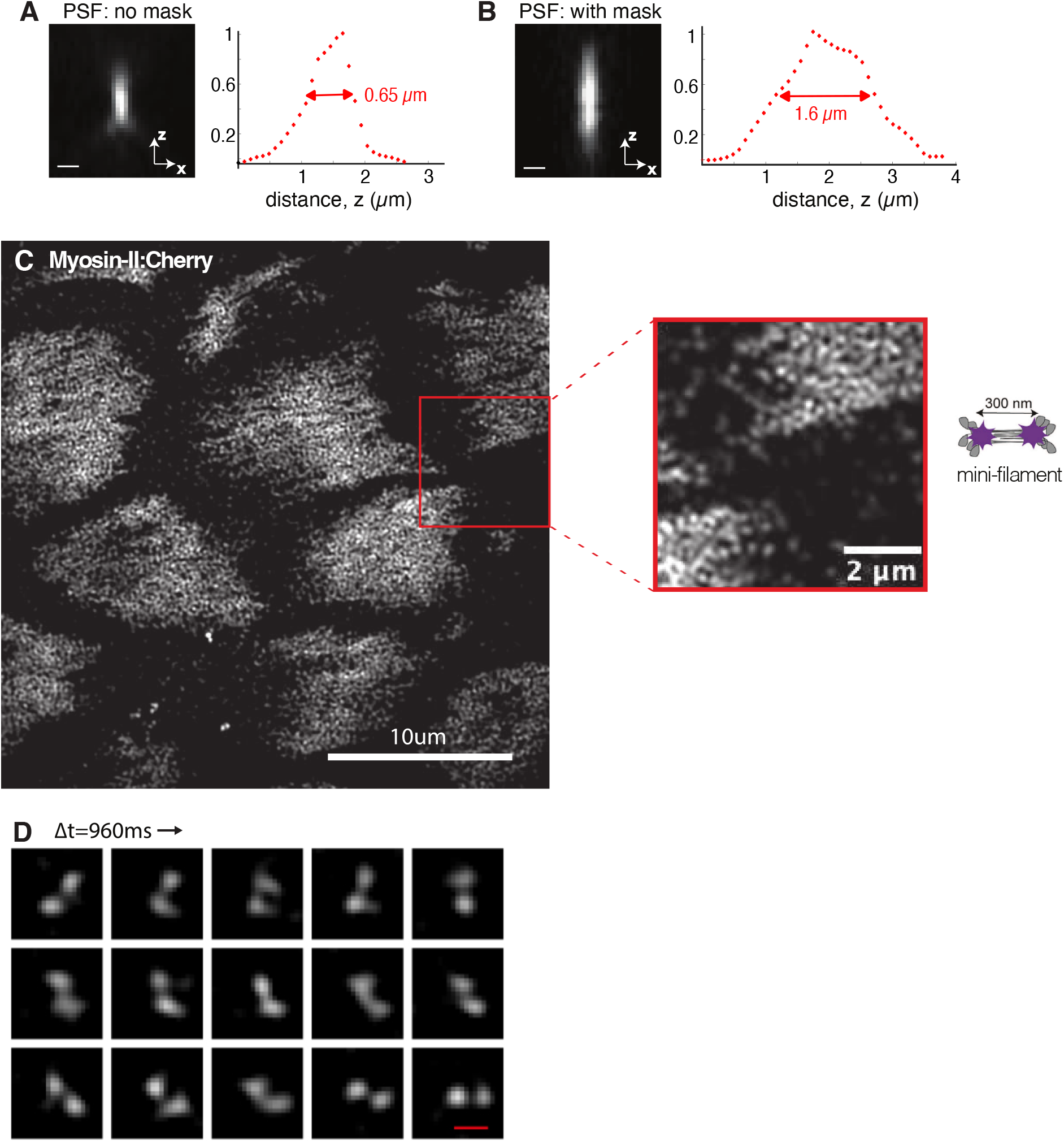
EDF-RIM using a stair-step phase mask. The ETL was replaced with a stair-step phase mask^37^. (A,B) The net result is an extension of the axial PSF from 0.65*µm* to 1.60*µm*. (C) Application to MyoII imaging in a *Drosophila* egg chamber. The schematic on the right shows the molecular structure of a typical mini-filament. The anti-polar arrangement leads to a fluorescent structure consisting of two fluorescent points, rigidly linked and spaced by approximately 300nm. (D) Close-up on sequences, imaged with a 960 ms interval, showing the rotation of an individual Myosin-II mini-filament (E), typical behavior that were described in^55^. Scales bars=300 nm when not specified.

**Figure SM1.**
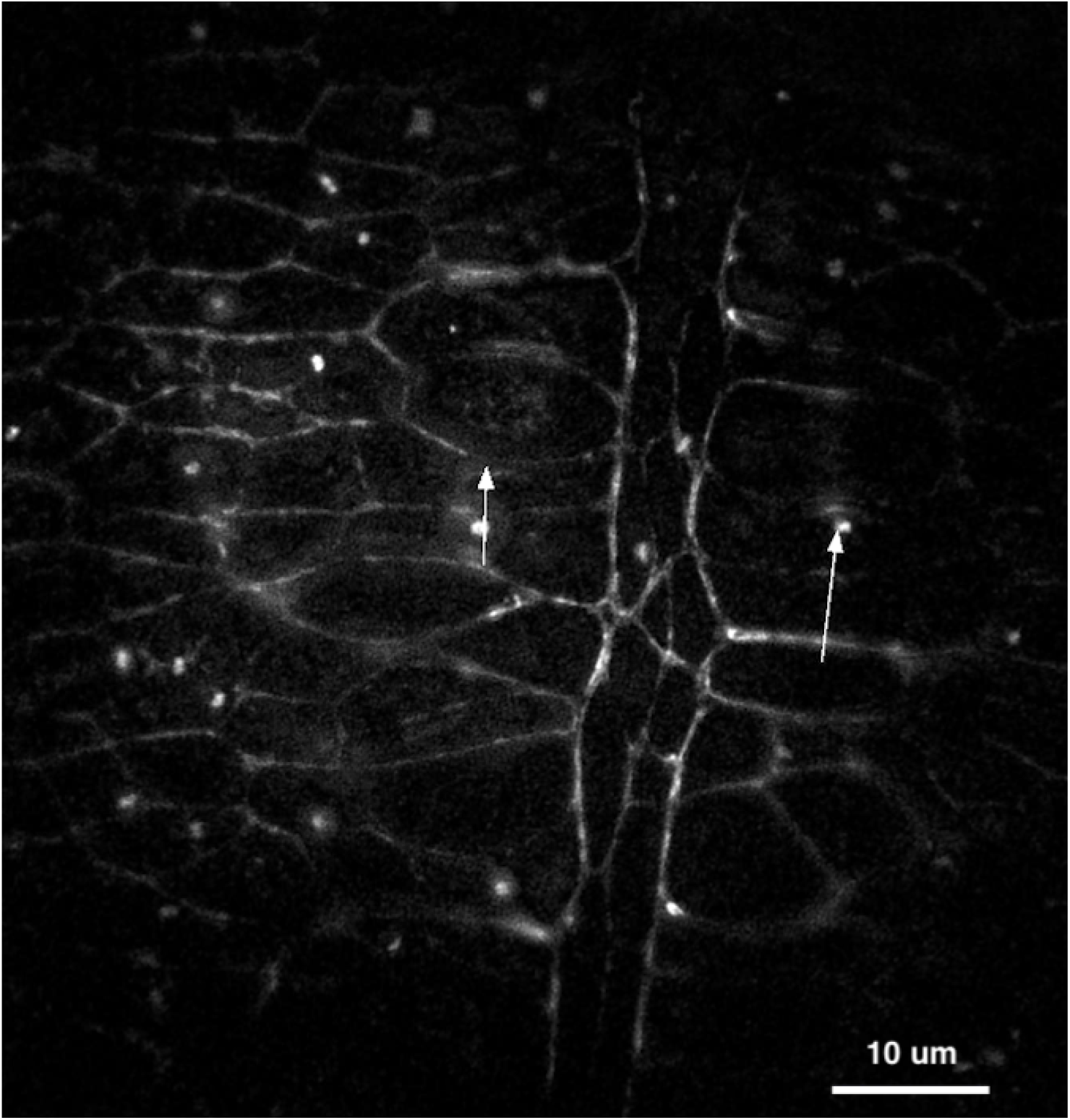
Supplementary movie 1 caption. Myosin:TagRFP imaging in the Drosophila embryo with EDF-RIM, in link with Fig. S3. White arrows point to closing cytokinetic rings. The temporal resolution is improved compared to Supplementary Movie 2.

**Figure SM2.**
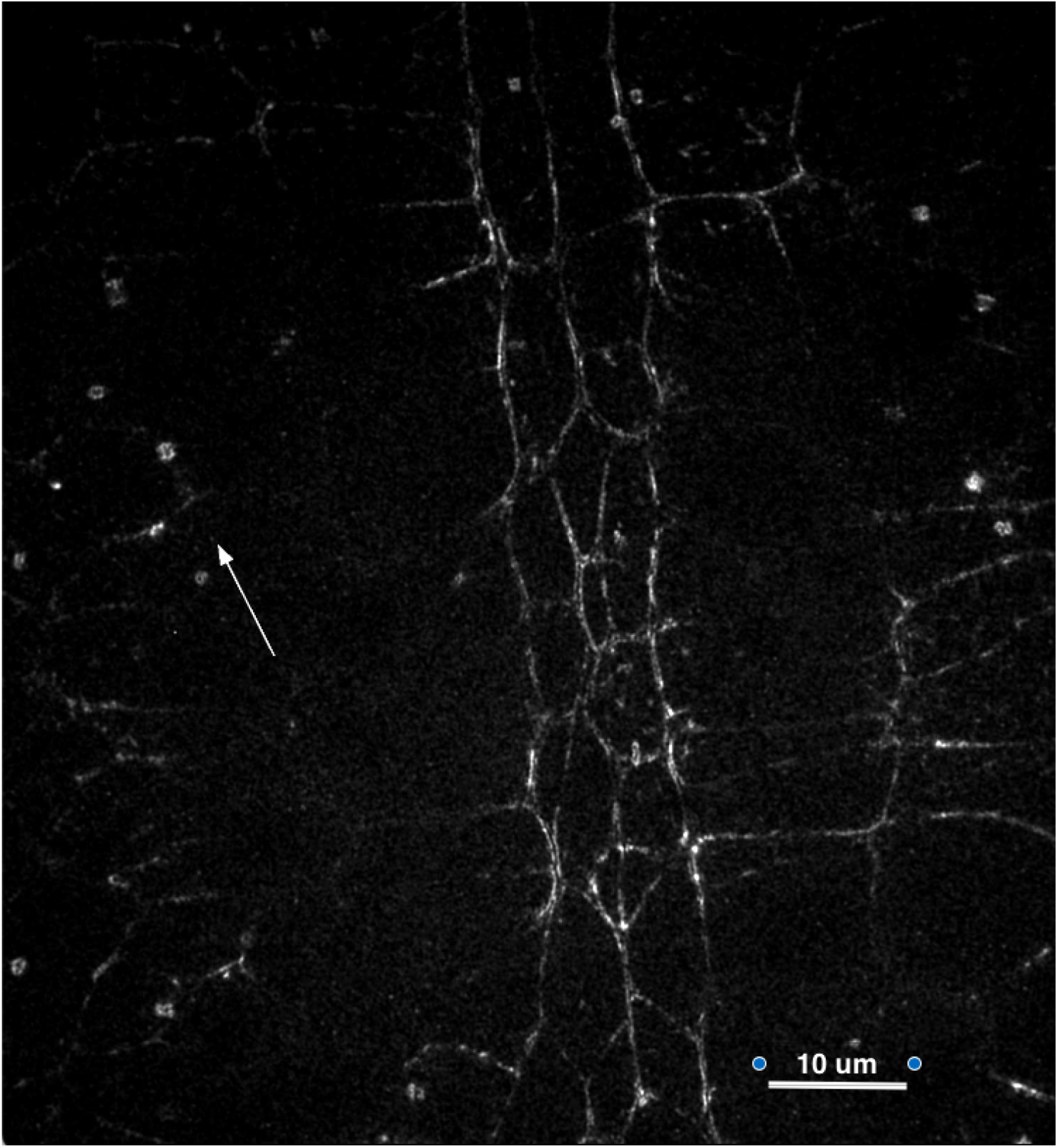
Supplementary movie 2 caption. Myosin:TagRFP imaging in the Drosophila embryo with 2D-RIM in link with Fig. S3. Multiple planes are acquired sequentially, and summed for projection. White arrows point to closing cytokinetic rings. The temporal resolution is decreased compared to Supplementary Movie 1.

**Figure SM3.**
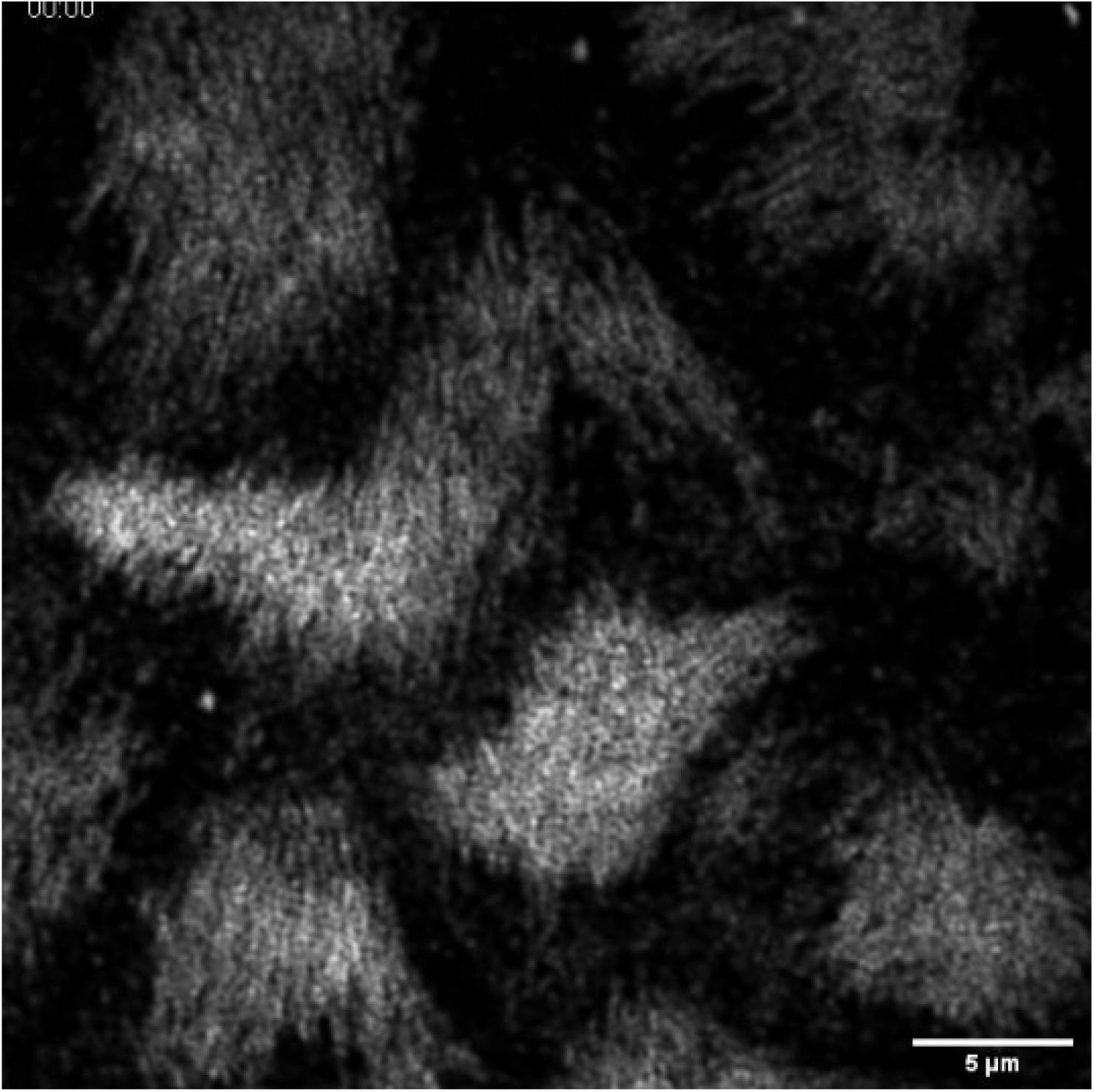
Supplementary movie 3 caption. Myosin-II:GFP in the basal surface of follicle cells from a Drosophila egg chamber in link with Fig. S5C.

**Figure SM4.**
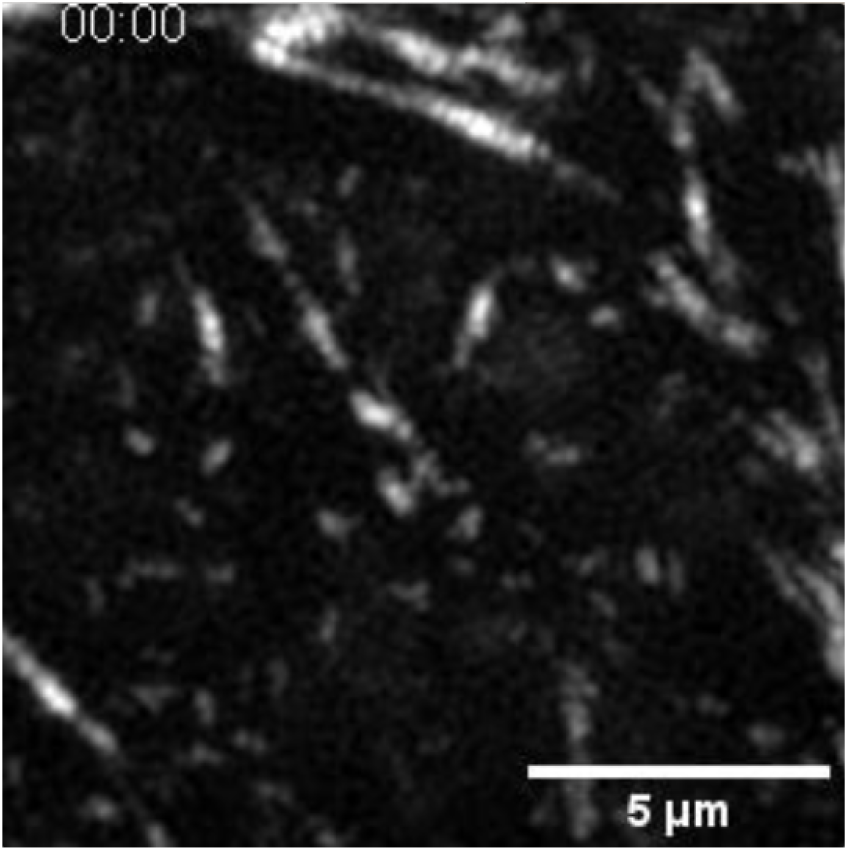
Supplementary movie 4 caption. Myosin-II:GFP in the basal surface of follicle cells from a Drosophila egg chamber in link with Fig. S5D

